# Bi-axial orientation could explain range expansion in a migratory songbird

**DOI:** 10.1101/2022.05.04.490589

**Authors:** Joe Wynn, Guillermo Fandos, Kira Delmore, Benjamin M. Van Doren, Thord Fransson, Miriam Liedvogel

## Abstract

The likelihood of a new migratory route evolving is a function of the associated fitness payoff, and the probability that the route arises in the first place. Cross-breeding studies suggest that young birds migrate in a direction intermediate between their parents, though this would seemingly not explain how highly divergent migratory trajectories arise in apparently sympatric populations. It has been suggested that diametrically opposed ‘reverse’ migratory trajectories might be surprisingly common, and if such routes were heritable it follows that they could underlie the rapid evolution of divergent migratory trajectories. Here, we used Eurasian blackcap (*Sylvia atricapilla*; ‘blackcap’) ringing recoveries and geolocator trajectories to investigate whether a recently-evolved northwards autumn migratory route could be explained by the reversal of each individual’s expected southwards migratory direction. We found that northwards migrants were recovered closer to the sites specified by a precise axis reversal than would be expected by chance, consistent with the rapid evolution of new migratory routes via bi-axial variation in orientation. We suggest that the surprisingly high probability of axis reversal might allow birds to expand their wintering ranges rapidly, and hence propose that understanding how direction is encoded is crucial when characterising the genetic basis of migratory direction and how this relates to route evolution.

## Introduction

Genetic inheritance of a migratory trajectory through 3-dimensional space appears extraordinary, yet is seemingly at the heart of how migratory routes might pass between generations. In addition to genetic inheritance, migratory phenotypes across avian taxa are also informed by asocial learning (Thorup et al., 2003; Wynn et al., 2020) and cultural inheritance (Chernetsov et al., 2004; Mueller et al., 2013). However, the contribution of genetic information to migratory direction would seem substantial in species moving long distances early in life unaccompanied by related conspecifics. In birds, inherited directional information is typically, though not universally (Thorup et al., 2020), assumed to comprise ‘clock and compass’ vector orientation: a compass to determine direction, and a clock to determine when to start and when to end migration (Berthold et al., 2013). This hypothesis is supported by experimental and observational studies, with naïve birds unable to compensate for displacement from the conventional migratory trajectory (Perdeck, 1958; Thorup et al., 2007) and following straight-line courses that accumulate error over time (consistent with vector navigation; (Mouritsen and Mouritsen, 2000; Wynn et al., 2021; Yoda et al., 2017).

Cross-breeding experiments between birds with different migratory routes suggest that additive genetic variance might inform migratory direction, with the progeny of birds with distinct migratory directions following trajectories that seemingly represent intermediate directions (Delmore and Irwin, 2014; Helbig, 1991). However, within such a model of genetic inheritance, it is difficult to account for the occurrence of highly divergent migratory directions. For example, small numbers of Eurasian blackcaps have been observed to increasingly migrate north in the autumn, with this shift linked to both changing climate and increasing availability of artificially provided food (Berthold et al., 1992; Plummer et al., 2015; Van Doren et al., 2021). North-migrating blackcaps are seemingly sympatric with south-migrating conspecifics (Delmore et al., 2020b), and hence it is difficult to reconcile such divergent migratory trajectories with classically considered additive genetic variance (given the apparent absence of any intermediate phenotypes). Assortative mating based on migratory destination has been reported (Bearhop et al., 2005), and whilst this might explain how divergent routes are maintained within a population this would not necessarily explain how they evolve in the first place.

It has been suggested that vagrant songbirds are disproportionately abundant in a direction precisely opposing the normal migratory direction, though such trends are difficult to verify beyond anecdote due to necessarily small sample sizes and/or biases in the distribution of observers (Gilroy and Lees, 2003; Thorup, 1998; Vinicombe, 1996). Whilst it is unclear what would cause markedly bi-axial variation in migratory populations, and indeed whether such variation would be heritable, this ‘reverse migration’ could nonetheless provide a mechanism by which highly divergent migratory routes evolve and persist within sympatric populations. It is, then, of interest whether (i) divergent migratory routes are quantifiably consistent with biaxial variation in orientation and (ii) if so, what might explain such variation. Here, we used ringing recoveries made over the last century (n = 78) alongside geolocator positions gathered in the years 2016-19 (n = 33) to investigate whether northwards migration by Eurasian blackcaps is likely to represent a reversal of the population expected migratory direction. Blackcaps have a well-documented bi-modal southwards migratory strategy, with birds from western Europe migrating south-west around the Mediterranean Sea and birds breeding in eastern Europe migrating south-east (with a narrow hybrid zone running north to south in the middle, in which intermediate directions are common; (Delmore et al., 2020b). We might expect, therefore, that northwards-migrating birds from either side of the migratory divide might ‘criss-cross’ in their wintering routes, with birds from south-east Europe migrating north-west and birds from south-west Europe migrating north-east.

## Results

To investigate this, we tested whether birds were recovered closer to the site predicted by a reversal of their anticipated southbound migratory route, than would be expected by chance. Using historic ringing records, we can define a likely southwards migratory route for each bird as the circular mean of all southwards migratory routes recorded in the vicinity (see Methods; see Figure S5), which in turn can be reversed to define the bearing predicted by an axis reversal. By taking this bearing, and moving the observed distance along it, we can specify an *a priori* expected recovery/wintering location under an axis-reversal hypothesis for each bird. Consequently, by comparing the Great Circle Distance from the observed recovery site to the recovery site expected under axis reversal (an ‘observed-versus-expected’ distance; see Methods; Wynn et al., 2022) we can measure how well independently collected ringing and geolocator data fit our hypothesis. The smaller this distance, the better the fit between the observed movements and those predicted by bi-axial variation.

To investigate whether these distances were smaller than would be expected by chance, we can compare the observed-versus-expected distances to those generated by equivalent, but random ‘null’ northwards migrations. The null model for each bird was constrained to move the same distance as the real bird, and be recovered at real recovery sites, but with the northwards movement direction selected at random from the population (see Methods, Wynn et al., 2022). This ensured that the null model took topography, habitat suitability and ringing effort into account, ensuring that biases in these factors did not lead to falsely positive results. In turn, by calculating how many times the null birds were closer to the expected recovery site than the observed birds, we could ascertain the probability that the observed data fit the hypothesis better than would be expected by chance (i.e. a p-value; see Figure 1 for a summary).

**Figure 1:**
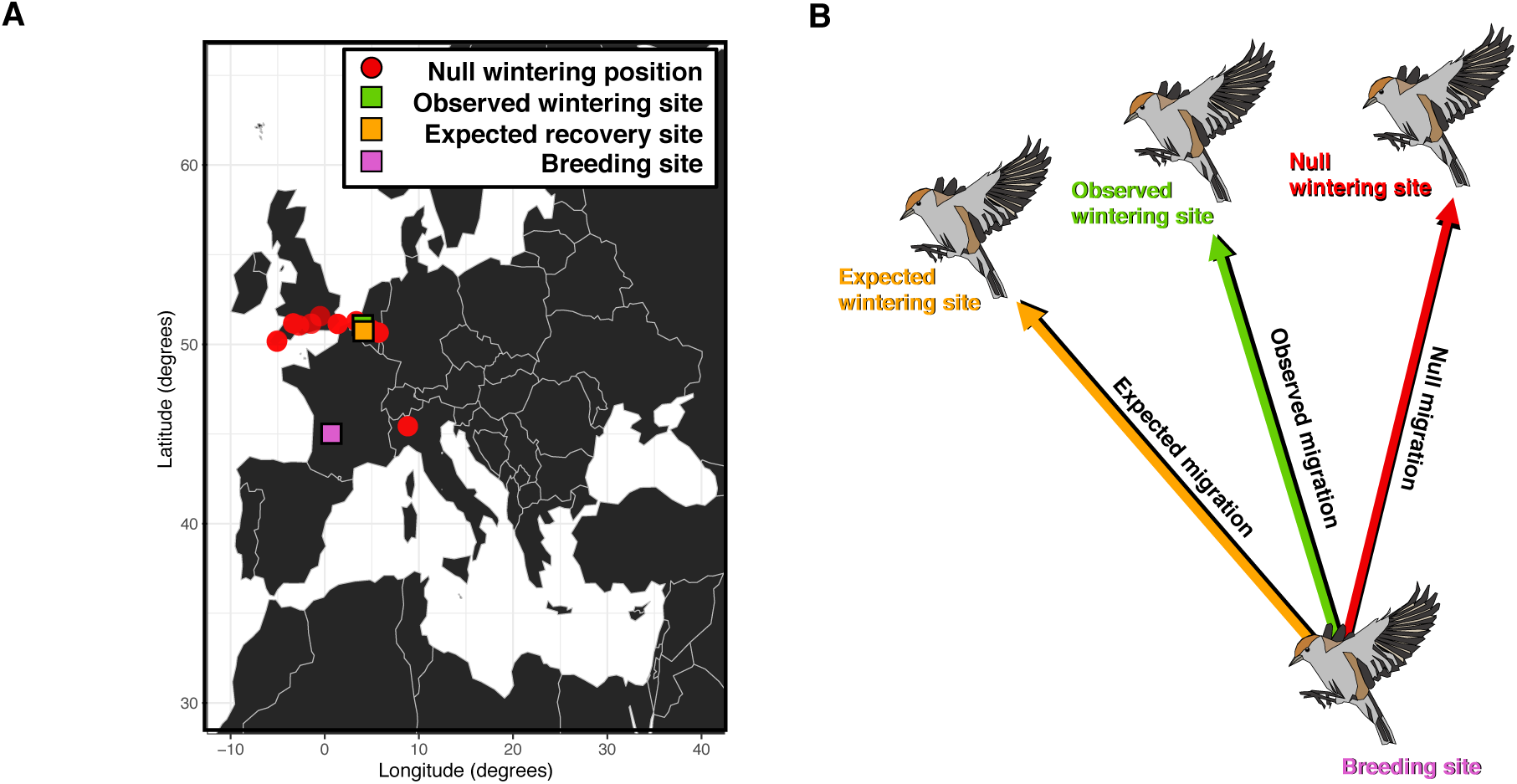
A null model of blackcap northwards migration. A) An example of how the distances between the expected wintering site and the observed/null wintering positions vary for a blackcap breeding in south-west France (purple). B) Schematic outlining the quantities used in our statistical analysis: the null migrations generated in our randomisation (red, see Methods), the observed recovery site (green) and the recovery site expected under a model of bi-axial directional variation (orange).

Using this randomisation, we found that blackcaps were recovered significantly closer to the site predicted by an axis reversal than would be expected owing to chance. This was true both when considering the mean distance between the observed and expected recovery sites (randomisation; p < 0.0001) and the median distance between the observed and expected recovery sites (randomisation; p < 0.0001). The fact that both the population mean and median observed-versus-expected distances were smaller than would be expected by chance suggests that variation in migratory direction is consistent with bi-axial orientation, even when biases caused by ringing effort, habitat distribution and topography are taken into account (see Figure 2). This would imply that (i) bi-axial orientational variance facilitates wintering range expansion in blackcaps and (ii) given that the pattern is unlikely to reflect the constraints imposed on blackcaps by topography and habitat, that this likely reflects bi-axial orientation variation in the genetically determined migratory direction.

**Figure 2:**
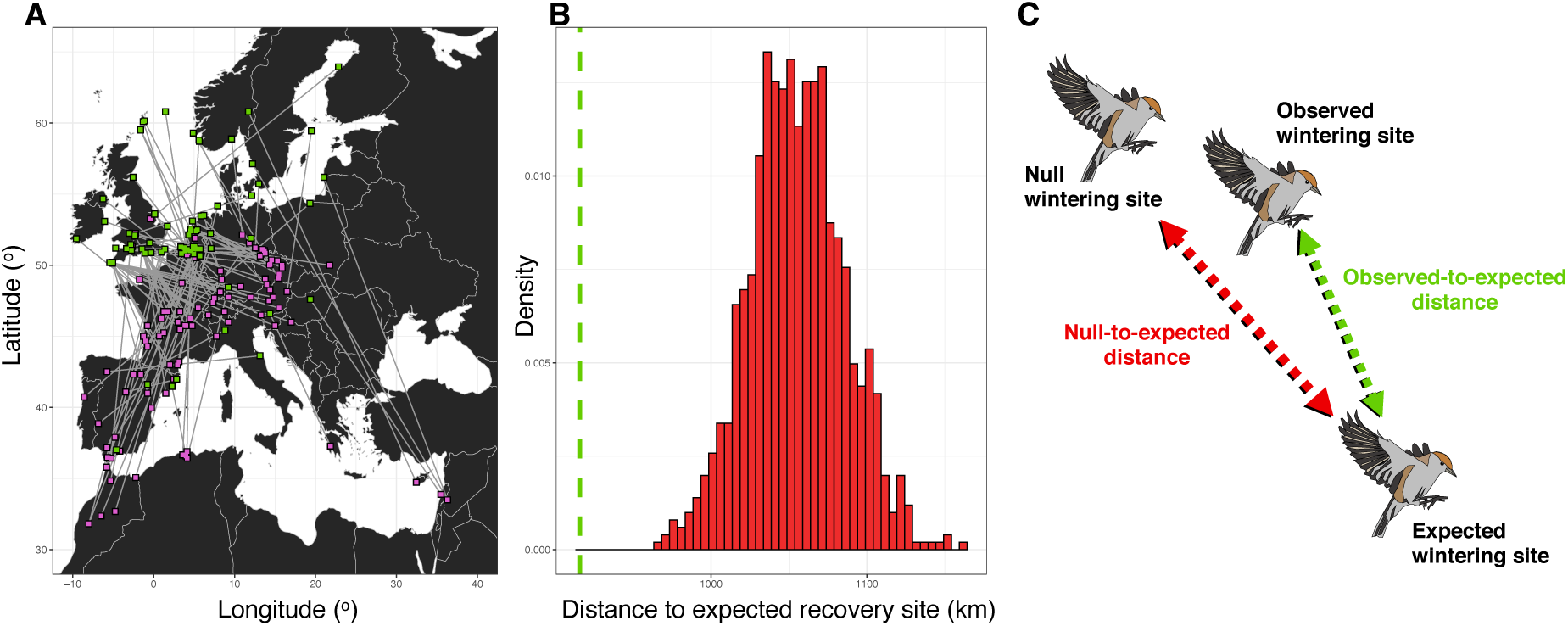
Axial variation in the migratory routes of Eurasian blackcaps. A) The breeding (pink) and wintering (green) sites of north-migrating blackcaps used in this analysis, with each recovery linked by a grey line between the breeding and non-breeding sites. B) A histogram showing the mean distance between null recovery sites and the expected recovery site, with the true observed-to-expected distance shown as a green dashed line. C) A schematic showing how the observed-versus-expected and null-versus-expected recovery sites are calculated for inclusion in the histogram.

## Discussion

Though it is apparent that additive genetic variance with progeny inheriting an intermediate direction between both parents would not predict the observed pattern, it isn’t clear what would. One possibility is that variance in bi-axial direction could be explained by variance in the ‘clock’ that regulates the time at which outbound and return migration trajectories are executed. This could be a similar phenomenon to the complete reversal of breeding and migratory phenology recently observed in South American barn swallows (*Hirundo rustica*; Areta et al., 2021), which in turn would suggest that axis reversal could represent the execution of the spring migratory trajectory in autumn. However, if this were the case it is unclear why breeding phenology would not simultaneously be reversed, as is observed in the barn swallow, and hence it is perhaps more likely that bi-axial orientation instead reflects the mechanism by which orientation information is encoded.

To discuss precisely how directional information might be encoded such that bi-axial directional variation occurs is speculative. Nonetheless, it is possible that these divergent migratory routes reflect the axial encoding of direction, with birds storing the autumn migratory route as an axis of direction alongside a preference for a given pole of that axis (Wynn et al., 2022). Indeed, the sensory and cognitive bases of avian compass orientation—where magnetic and star compass orientation is conducted relative to the pole/equator (and thus is symmetrical in both directions) rather than north/south (i.e. directional)—makes this two-step process quite likely (Emlen, 1967a, b; Wiltschko and Wiltschko, 1972). Therefore, bi-axial orientational encoding might be surprisingly plausible. Such a mechanism would likely be an example of epistasis, with the effect of gene(s) coding for the migratory axis modulated by genes coding for directional preference along that axis. Bi-axiality in orientation would, then, arise as a deviant phenotype based on this interaction. In a more general sense, epistatic interactions between genes at difference loci might produce bi-axial orientation through another mechanism, though, no extant hypothesis outlines what this mechanism might be, and indeed examples of epistatic interactions are rare in animal behaviour (e.g. Godoy-Herrera et al., 2004; Yamamoto et al., 2009). Hence, it is apparent that further research is essential when considering how genetic differences correspond to variation in migratory direction.

Further, understanding the causes of bi-axiality in migratory direction is of some importance when investigating which gene(s) or regulatory region(s) is/are responsible for determining migratory direction. Recent advances in the affordability and resolution of biologging and genetic sequencing technology make genome-wide associations with migratory phenotype possible (e.g. Delmore et al., 2020a; Delmore et al., 2016; Toews et al., 2019), and if such techniques were applied to migratory direction it is key that the mechanism by which directional information is encoded is understood fully. Without a complete understanding of how direction varies, and is in turn encoded, such analyses might fail to capture variance in migratory direction correctly, and hence lead to the erroneous association/disassociation of loci with directional traits. For example, if blackcap direction was indeed stored – first – as a migratory axis and then – second – as a directional preference for a given pole of that axis, we would expect that birds migrating south-west should be genetically more similar to birds that migrate north-east than birds migrating due south at loci associated with migratory direction. Investigating which genes/genomic regions specifically moderate migratory direction is, then, contingent on correctly understanding the means by which direction is encoded.

Whilst it is necessarily difficult to draw conclusions based solely upon correlative analyses, we nonetheless believe our results inform upon both (i) the mechanisms by which migratory directions are encoded and (ii) how future studies into the genetic correlates of migration direction might be conducted. Further, our results make predictions as to where novel migratory routes might evolve into the future, and might then be of some utility when considered within the context of recent environmental shifts (e.g. changes in land use and climate). Therefore, we suggest that further studies utilising historic ringing recoveries or tracking data could be of considerable interest when considering how long-term ecological changes might interact with the mechanisms governing migratory direction to produce novel migratory routes.

## Methods

### Selecting ringing records

In this manuscript we sought to investigate whether the wintering positions of autumn north-migrating blackcaps, inferred from historic ringing records and geolocator trajectories, were likely to reflect a reversal of ‘normal’ southward migratory direction in autumn, and hence inform on whether migratory direction is more likely to vary in a bi-axial manner than would be expected by chance. Given the spatiotemporal biases associated with ‘citizen science’ data, and the sensitivity of any analysis of migratory direction to these biases, it is therefore key to subset ringing data to include only instances in which the same bird is caught during both the breeding and the non-breeding periods. From these birds, we can then select individuals where the non-breeding position is located north of the breeding site. In previous studies that utilise ringing records to study navigation, the problem of how to categorise birds as breeding was solved by exhaustively subsetting the data using different criteria to ensure that the method of subsetting didn’t drive the results (Paradis et al., 1998; Wynn et al., 2022). Here, however, we sought to improve upon this method by including phenotypic markers of breeding and migration to ensure that birds recaptured on migration were effectively excluded from the analysis.

To remove migrating birds, we sought to ascertain the point in time at which breeding became far more likely than migration. The onset of breeding in blackcaps is characterised by the development of both a brood patch in females and a cloacal protrusion in males, whilst very young blackcaps that have not completed post-juvenile moult (and hence haven’t left the breeding site) have a field-identifiable plumage (EURING code ‘3J’). Similarly, birds ringed as chicks in the nest are recorded as such in EURING. In contrast, migrating blackcaps typically have a higher fuel load than breeding birds (Svensson, 1970), which manifests as subcutaneous fat. Since both the breeding and migratory phenotypes are distinctly recognisable at the point of ringing, we first sought to use ringing data to isolate the time of year at which breeding became more likely than migrating. We did this by comparing the probability of occurrence of breeding versus non-breeding phenotypes at different times of year, defining breeding birds as either birds ringed as chicks in the nest or free-flying birds with a cloacal protuberance, brood patch or juvenile plumage, and migrant birds as having a fat score of > 3 (meaning a high degree of visible subcutaneous fat) or the equivalent fuel load of 17.5% as calculated using mass and maximum wing chord based on the methods outlined in (Kelsey et al., 2019).

We divided Europe up into a 5° x 5° grid, and for each grid point we subsetted the overall EURING database for every bird with a breeding phenotype within 5° of the grid point in question. We then repeated this process for both spring and autumn migratory phenotypes. If the sample size for each of spring migration, breeding and autumn migration was > 10, we then calculated a density curve (bandwidth = 10) of the recorded Julian dates for spring migration, breeding and autumn migration (see Figure S1). The points at which spring migration density < breeding density (i.e. the point at which a ringing event was more likely to represent a breeding bird than a migrant) and the point at which breeding density < autumn migration density (i.e. the point at which a ringing event was more likely to represent a migrant bird than a breeding bird) were then isolated for each grid square. Based on this categorisation, we could then identify the respective dates at which breeding was likely to commence (see Figure S2) and end (see Figure S3) for all points in Europe where ringing data were available.

Whilst it would be possible to use the calculated start and end of breeding from each point in Europe where data exists to subset our data to include only birds likely to be breeding, we chose not to do this, since a) this would lead to a paucity of data from ringing schemes where biometric information is not readily reported, and b) just because migration has become less likely than breeding this doesn’t mean migration isn’t occurring. As such, we sought to define breeding in our analysis as either birds recorded with a breeding phenotype as defined above, or recorded from within a core breeding window of 15^th^ June – 15^th^ July in our analysis. Within this time window birds from all over Europe were very unlikely to be on migration (see Figures S2 and S3), and the window used was consistent with the breeding phenology and migratory timings reported in previous studies (Delmore et al., 2020b). The position of this window relative to overall migratory and breeding phenology from across Europe is shown in figure S4. Only birds that showed northwards autumn migratory direction and moved > 500km between ringing and recovery (n = 78) were retained in our main analysis.

In our analysis we also included wintering/breeding positions as estimated from geolocator (GLS) tracked blackcaps. When considering geolocator trajectories (n = 33), breeding and wintering positions were ascertained using the geolight package (Lisovski and Hahn, 2012) using the methods outlined in (Delmore et al., 2020b).

### Randomisation analysis

For each north-migrating bird we attempted to quantify how close the bird was recovered to the site expected by an axis reversal. To estimate an expected southwards migratory direction for a population from which a given north-migrating bird originates, we calculated the (circular) mean southwards migratory direction taken by the 50 closest southwards breedingto-wintering ringing records (see above) to the focal north-migrating bird (see Figure S5). By adding 180& to this direction, we could then estimate the northwards migratory trajectory expected under an axis-reversal. By moving the observed distance along this trajectory, we could, therefore, estimate the site at which a bird might be expected to be recovered, and by measuring the distance between the observed recovery site and this expected recovery site we could estimate how well the observed trajectories fit our theory. The smaller this observedversus-expected distance is, the better the fit of the model.

If a bi-axial variation hypothesis is correct, we would anticipate that this observed-versus-expected distance to be smaller than would be expected by chance. However, it is extremely difficult to interpret these distances bald since we have no realistic *a priori* expectation as to what these distances should be if birds migrated north at random. As a consequence, it is key to establish a realistic null model of northwards migration that does not directly stem from axis reversal. If the observed birds are closer to the expected recovery site than the simulated null model birds are, this would support the idea that northwards movement of blackcaps is driven by variation in an inherited bi-axial migratory program.

To create realistic null wintering sites for each bird, we first assigned a northward bearing selected at random (with replacement) from all recorded northward bearings. Second, we isolated the site that was the same distance along the randomly selected bearing as the observed bird moved. Finally, we then constrained the null recovery site to the nearest site where a bird was ringed, so as to ensure that the null birds had the same distribution constraints as real birds. Importantly, ringed birds were constrained to take randomly selected migratory directions and be recovered at wintering sites inherited from other ringed birds, and similarly the null trajectories simulated for GLS-tracked birds were derived from other GLS tracked birds. This ensured that the constraints on wintering sites seen in the observed birds were carried forward into the null model.

We ran our null model 100,000 times, each time calculating a mean and a median observedversus-expected distance, which we could compare to our empirical mean and median observed-versus-expected distances. We then calculated the number of times the null model had a smaller observed-versus-expected distance than was observed empirically, and hence calculated a p-value for both the mean and median observed-versus-expected distances. All p-values were corrected for the 2-tailed tests performed by multiplying the p-value by 2.

## Acknowledgements

This work was supported through funding from the Max Planck Society (MFFALIMN0001) and the DFG (SFB 1372 – Magnetoreception and Navigation in Vertebrates). We’d like to thank both the European Union for Bird Ringing (EURING) data bank for compiling the data used in this study and everyone in Europe who’s ringed a blackcap, without whom studies like this are not possible. We’d also like to thank Oliver Padget, Tim Guilford, Paris Jaggers, Joe Morford and Katrina Davies for comments on an early version of this concept.

## Author contributions

Analysis was conducted by J.W. with input from G.F. and M.L. The manuscript was written by J.W., with input from M.L., and was revised with input from K.D. and B.V.D. GLS data presented was gathered by K.D., B.V.D. and M.L.

## Supplementary material

### Analysis with the southernmost breeding birds removed

We re-ran our randomisation analysis considering only birds that were recorded breeding at a latitude greater than 35°. We did this since even though birds found below this latitude were retained using our objective subsetting methods, these movements seem surprising given the known ecology of blackcaps. These included birds apparently breeding in North Africa and wintering in the UK, and those breeding in Cyprus and Greece and wintering in Northern Europe. As such, we sought to ensure that such atypical records were not driving our results. We found that both the median (randomisation; p < 0.0001) and mean (randomisation; p < 0.0001) distance between the site predicted by an axis reversal and the observed site was smaller than would be expected by chance (see Figure S5).

---

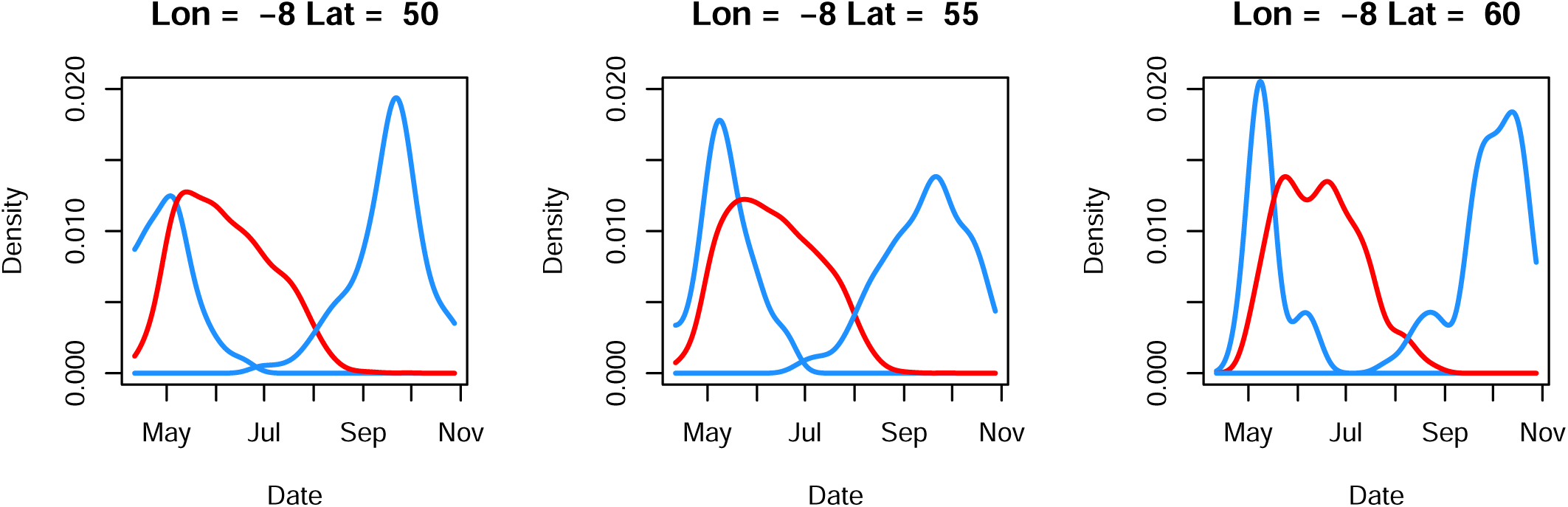

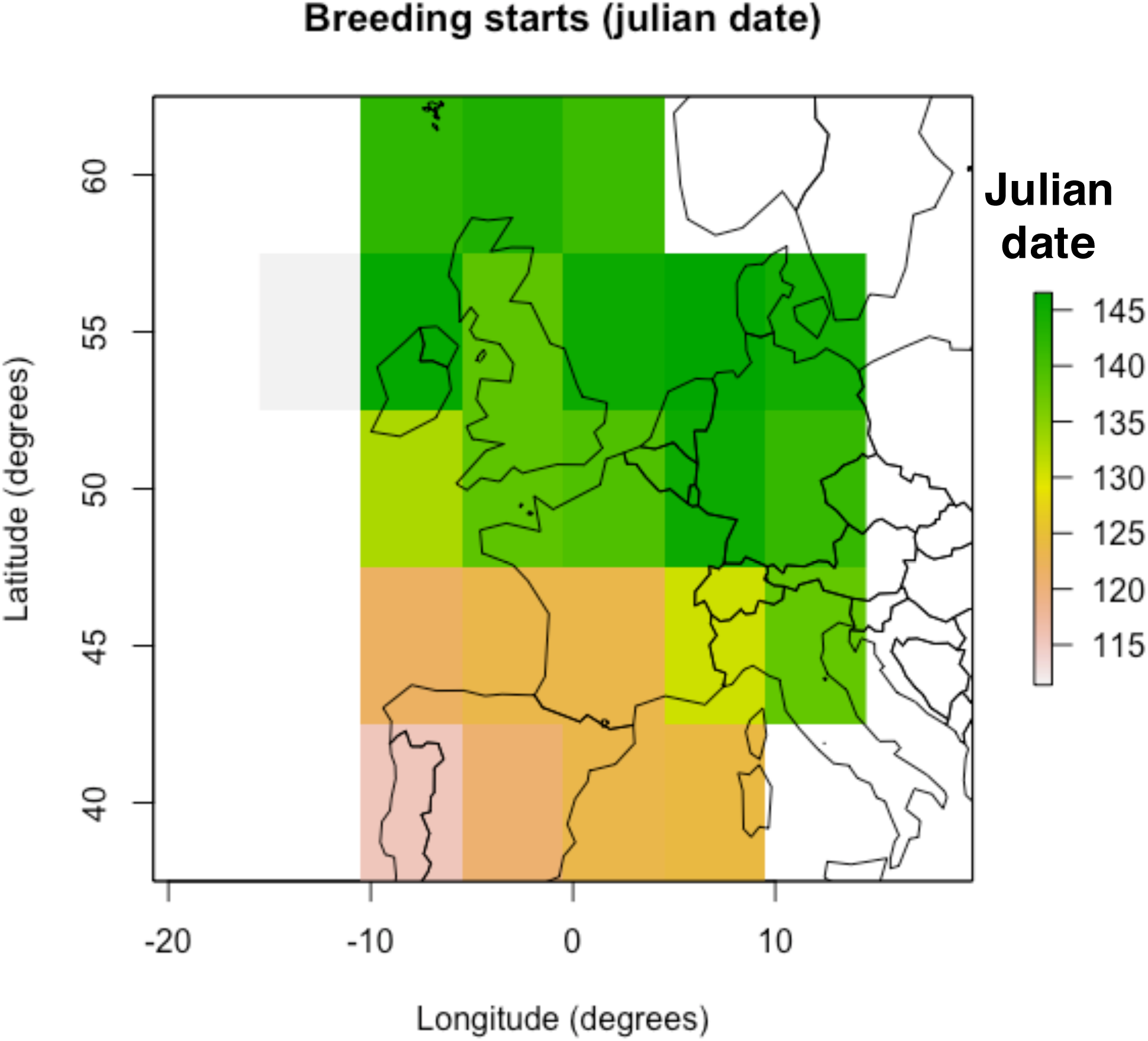

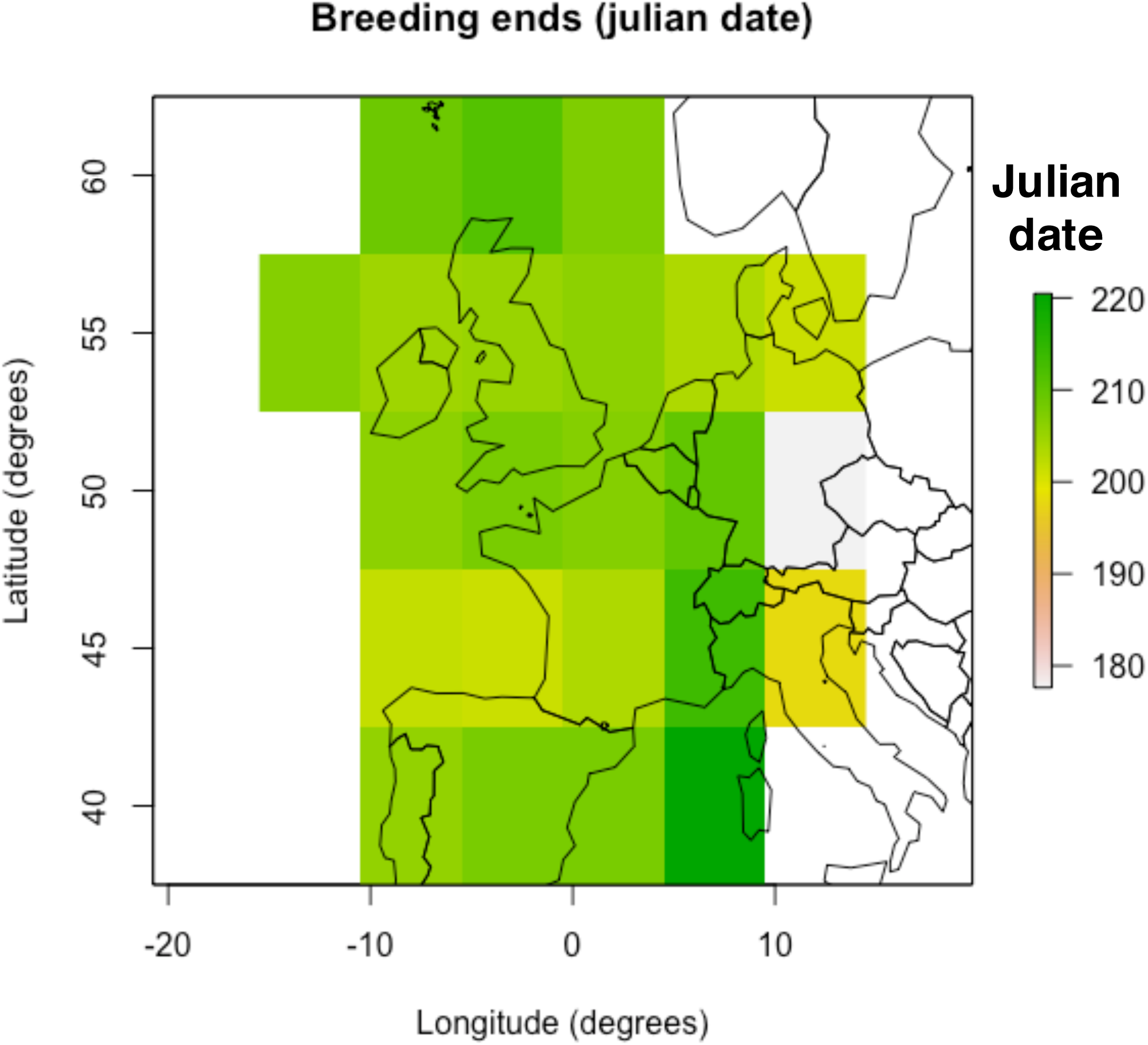

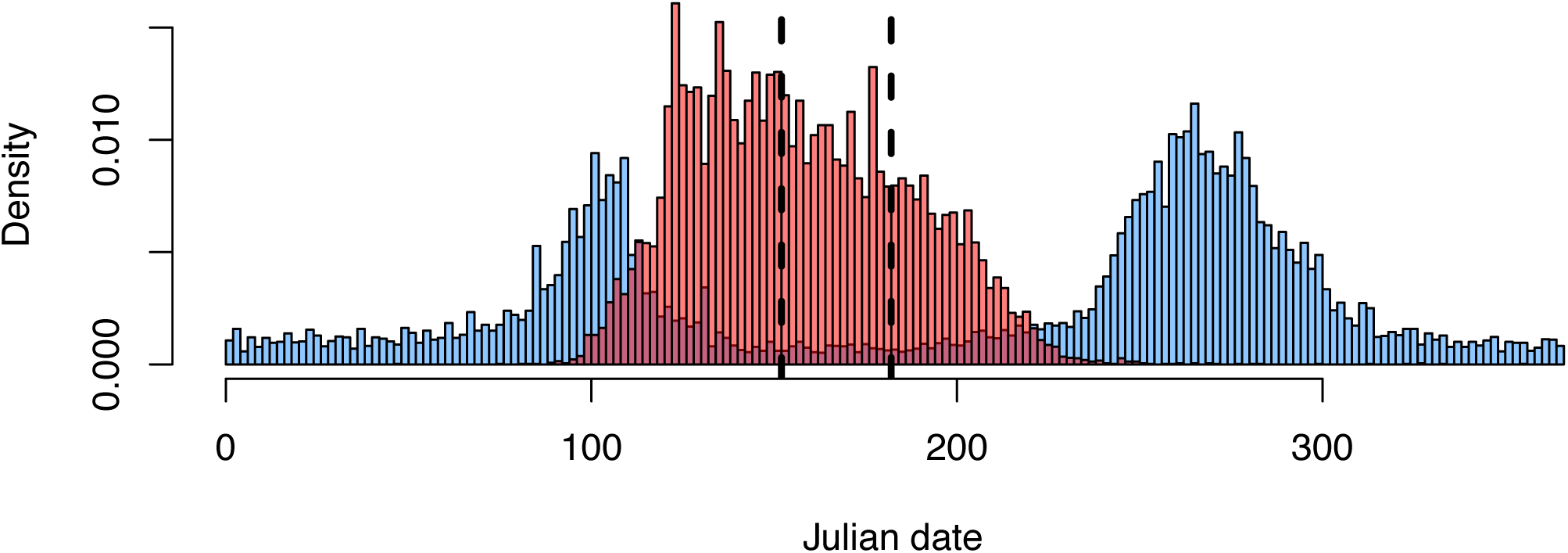

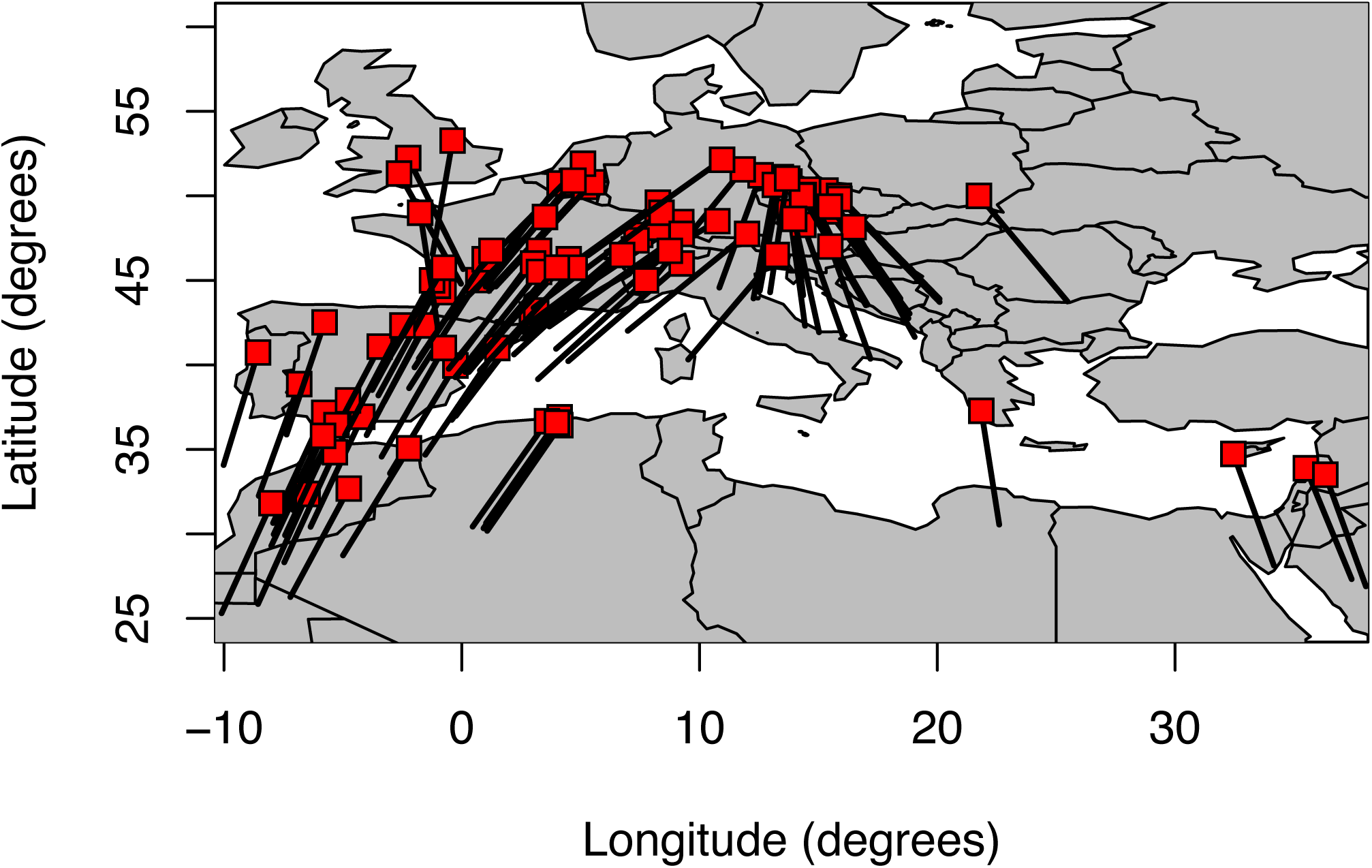

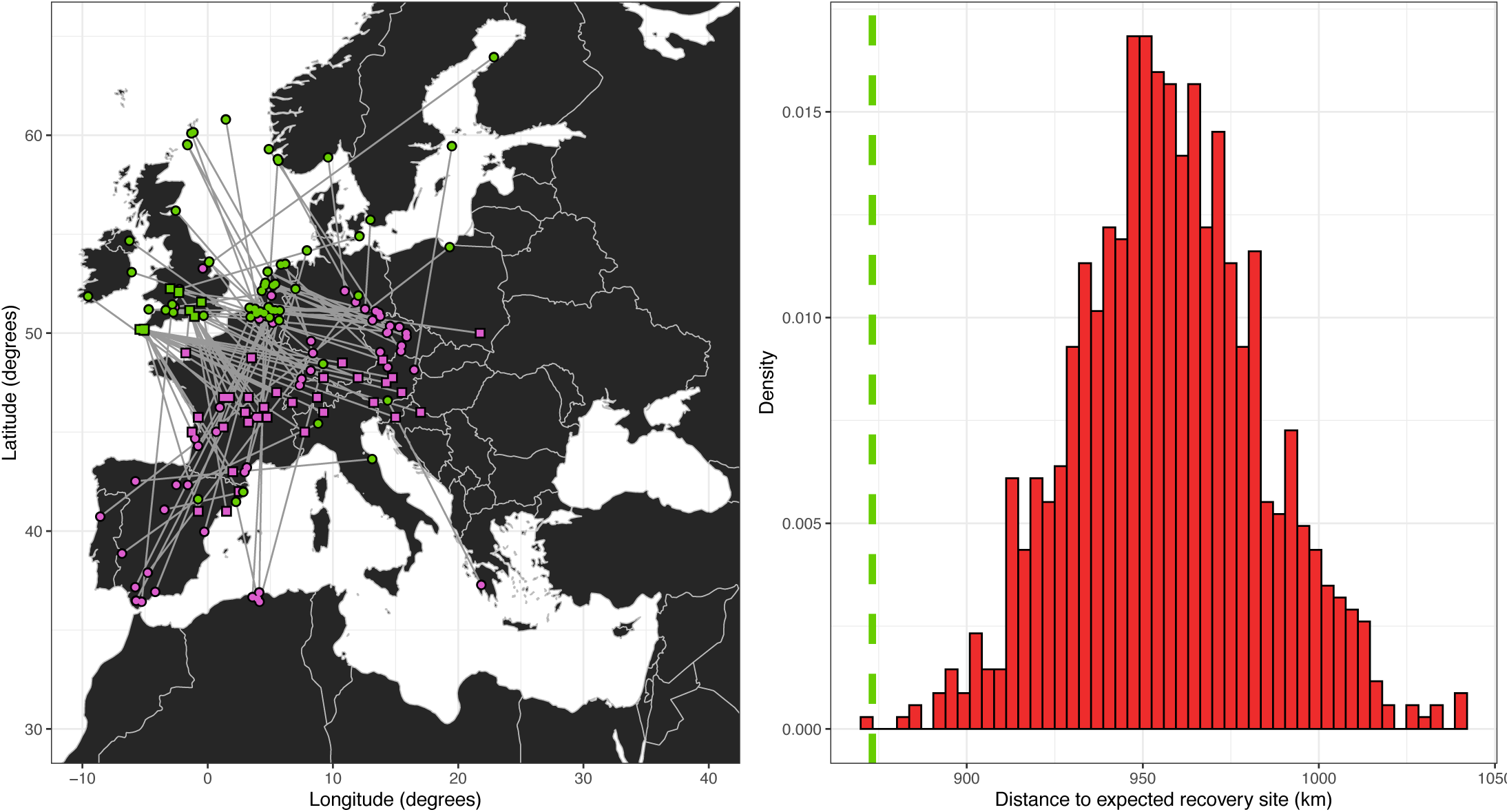

## References

Areta, J. I., S. A. Salvador, F. A. Gandoy, E. S. Bridge, F. C. Gorleri, T. M. Pegan, E. R. Gulson-Castillo, K. A. Hobson, and D. W. Winkler, 2021, Rapid adjustments of migration and life history in hemisphere-switching cliff swallows: Current Biology, v. 31, p. 2914-+.

Bearhop, S., W. Fiedler, R. W. Furness, S. C. Votier, S. Waldron, J. Newton, G. J. Bowen, P. Berthold, and K. Farnsworth, 2005, Assortative mating as a mechanism for rapid evolution of a migratory divide: Science, v. 310, p. 502–504.

Berthold, P., E. Gwinner, and E. Sonnenschein, 2013, Avian migration, Springer Science & Business Media.

Berthold, P., A. J. Helbig, G. Mohr, and U. Querner, 1992, Rapid microevolution of migratory behaviour in a wild bird species: Nature, v. 360, p. 668–670.

Chernetsov, N., P. Berthold, and U. Querner, 2004, Migratory orientation of first-year white storks (Ciconia ciconia): inherited information and social interactions: Journal of Experimental Biology, v. 207, p. 937–943.

Delmore, K., J. C. Illera, J. Perez-Tris, G. Segelbacher, J. S. L. Ramos, G. Durieux, J. Ishigohoka, and M. Liedvogel, 2020a, The evolutionary history and genomics of European blackcap migration: Elife, v. 9.

Delmore, K. E., and D. E. Irwin, 2014, Hybrid songbirds employ intermediate routes in a migratory divide: Ecology Letters, v. 17, p. 1211–1218.

Delmore, K. E., D. P. L. Toews, R. R. Germain, G. L. Owens, and D. E. Irwin, 2016, The Genetics of Seasonal Migration and Plumage Color: Current Biology, v. 26, p. 2167–2173.

Delmore, K. E., B. M. Van Doren, G. J. Conway, T. Curk, T. Garrido-Garduno, R. R. Germain, T. Hasselmann, D. Hiemer, H. P. van der Jeugd, H. Justen, J. S. L. Ramos, I. Maggini, B. S. Meyer, R. J. Phillips, M. Remisiewicz, G. C. M. Roberts, B. C. Sheldon, W. Vogl, and M. Liedvogel, 2020b, Individual variability and versatility in an eco-evolutionary model of avian migration: Proceedings of the Royal Society B-Biological Sciences, v. 287.

Emlen, S. T., 1967a, Migratory orientation in the Indigo Bunting, Passerina cyanea. Part II: Mechanism of celestial orientation: The Auk, v. 84, p. 463–489.

Emlen, S. T., 1967b, Migratory orientation in the indigo bunting, passerina cyanea: part i: evidence for use of celestial cues: The Auk, v. 84, p. 309–342.

Gilroy, J. J., and A. C. Lees, 2003, Vagrancy theories: are autumn vagrants really reverse migrants?: British Birds, v. 96, p. 427–438.

Godoy-Herrera, R., B. Burnet, and K. Connolly, 2004, Conservation and divergence of the genetic structure of larval foraging behaviour in two species of the Drosophila simulans clade: Heredity, v. 92, p. 14–19.

Helbig, A. J., 1991, Inheritance of migratory direction in a bird species: a cross-breeding experiment with SE-and SW-migrating blackcaps (Sylvia atricapilla): Behavioral Ecology and Sociobiology, v. 28, p. 9–12.

Kelsey, N. A., H. Schmaljohann, and F. Bairlein, 2019, A handy way to estimate lean body mass and fuel load from wing length: a quantitative approach using magnetic resonance data: Ringing & Migration, v. 34, p. 8–24.

Lisovski, S., and S. Hahn, 2012, GeoLight-processing and analysing light-based geolocator data in R: Methods in Ecology and Evolution, v. 3, p. 1055–1059.

Mouritsen, H., and O. Mouritsen, 2000, A mathematical expectation model for bird navigation based on the clock-and-compass strategy: Journal of Theoretical Biology, v. 207, p. 283–291.

Mueller, T., R. B. O’Hara, S. J. Converse, R. P. Urbanek, and W. F. Fagan, 2013, Social Learning of Migratory Performance: Science, v. 341, p. 999–1002.

Paradis, E., S. R. Baillie, W. J. Sutherland, and R. D. Gregory, 1998, Patterns of natal and breeding dispersal in birds: Journal of Animal Ecology, v. 67, p. 518–536.

Perdeck, A., 1958, Two types of orientation in migrating starlings, Sturnus yulgaris L., and Chaffinches, Fringilla coelebs L., as Revealed by Displacement Experiments: Ardea, v. 55, p. 1–3.

Plummer, K. E., G. M. Siriwardena, G. J. Conway, K. Risely, and M. P. Toms, 2015, Is supplementary feeding in gardens a driver of evolutionary change in a migratory bird species?: Global Change Biology, v. 21, p. 4353–4363.

Svensson, L., 1970, Identification guide to European passerines: Stockholm, Naturhistoriska Riksmuseet, 152 p. p.

Thorup, K., 1998, Vagrancy of Yellow-browed Warbler Phylloscopus inornatus and Pallas’s Warbler Ph. proregulus in north-west Europe: Misorientation on great circles?: Ringing & Migration, v. 19, p. 7–12.

Thorup, K., T. Alerstam, M. Hake, and N. Kjellen, 2003, Bird orientation: compensation for wind drift in migrating raptors is age dependent: Proceedings of the Royal Society B-Biological Sciences, v. 270, p. S8–S11.

Thorup, K., I. A. Bisson, M. S. Bowlin, R. A. Holland, J. C. Wingfield, M. Ramenofsky, and M. Wikelski, 2007, Evidence for a navigational map stretching across the continental US in a migratory songbird: Proceedings of the National Academy of Sciences of the United States of America, v. 104, p. 18115–18119.

Thorup, K., M. L. Vega, K. R. S. Snell, R. Lubkovskaia, M. Willemoes, S. Sjöberg, L. V. Sokolov, and V. Bulyuk, 2020, flying on their own wings: young and adult cuckoos respond similarly to long-distance displacement during migration: Scientific Reports, v. 10, p. 1–8.

Toews, D. P. L., S. A. Taylor, H. M. Streby, G. R. Kramer, and I. J. Lovette, 2019, Selection on VPS13A linked to migration in a songbird: Proceedings of the National Academy of Sciences of the United States of America, v. 116, p. 18272–18274.

Van Doren, B. M., G. J. Conway, R. J. Phillips, G. C. Evans, G. C. M. Roberts, M. Liedvogel, and B. Sheldon, 2021, Human activity shapes the wintering ecology of a migratory bird: Global Change Biology, v. 27, p. 2715–2727.

Vinicombe, K., 1996, Rare birds in Britain & Ireland : a photographic record: London, HarperCollins.

Wiltschko, W., and R. Wiltschko, 1972, Magnetic compass of European robins: Science, v. 176, p. 62–64.

Wynn, J., J. Collet, A. Prudor, A. Corbeau, O. Padget, T. Guilford, and H. Weimerskirch, 2020, Young frigatebirds learn how to compensate for wind drift: Proceedings of the Royal Society B-Biological Sciences, v. 287.

Wynn, J., T. Guilford, O. Padget, C. M. Perrins, N. McKee, N. Gillies, C. Tyson, B. Dean, H. Kirk, and A. L. Fayet, 2021, Early-life development of contrasting outbound and return migration routes in a long-lived seabird: Ibis.

Wynn, J., O. Padget, H. Mouritsen, J. Morford, P. Jaggers, and T. Guilford, 2022, Magnetic stop signs signal a European songbird’s arrival at the breeding site after migration: Science, v. 375, p. 446–449.

Yamamoto, A., R. R. H. Anholt, and T. F. C. Mackay, 2009, Epistatic interactions attenuate mutations affecting startle behaviour in Drosophila melanogaster: Genetics Research, v. 91, p. 373–382.

Yoda, K., T. Yamamoto, H. Suzuki, S. Matsumoto, M. Muller, and M. Yamamoto, 2017, Compass orientation drives naive pelagic seabirds to cross mountain ranges: Current Biology, v. 27, p. R1152–R1153.

